# Differential outcomes of infection by wild-type SARS-CoV-2 and the B.1.617.2 and B.1.1.529 variants of concern in K18-hACE2 transgenic mice

**DOI:** 10.1101/2023.09.08.556906

**Authors:** Yicheng He, Jill Henley, Philip Sell, Lucio Comai

**Affiliations:** Department of Molecular Microbiology and Immunology; Hastings Foundation and Wright Foundation BSL3 laboratory, Keck School of Medicine, University of Southern California, Los Angeles, CA 90089

**Keywords:** SARS-CoV-2, Delta and Omicron variants, infection, brain, cytokines, chemokines

## Abstract

**Background:** SARS-CoV-2 is a respiratory virus with neurological complications including loss of smell and taste, headache, and confusion that can persist for months or longer. Severe neuronal cell damage has also been reported in some cases. The objective of this study was to compare the infectivity of Wild-type virus, Delta and Omicron variants in transgenic mice that express the human angiotensin-converting enzyme 2 (hACE2) receptor under the control of the keratin 18 promoter (K18) and characterize the progression of infection and inflammatory response in the lung, brain medulla oblongata and olfactory bulbs of these animals. We hypothesized that Wild-type, Delta and Omicron differentially infect K18-hACE2 mice, thereby inducing distinct cellular responses.

**Methods:** K18-hACE2 female mice were intranasally infected with Wild-type, Delta, or Omicron variants and euthanized either at 3 days post-infection (dpi) or at the humane endpoint. None of the animals infected with the Omicron variant reached the humane endpoint and were euthanized at day 8 dpi. Virological and immunological analyses were performed in the lungs, olfactory bulbs, medulla oblongata, and brains.

**Results:** Mice infected with Wild-type and Delta display higher levels of viral RNA in the lungs than mice infected with Omicron at 3dpi. Viral RNA levels in the brains of mice infected with the Wild-type virus were however significantly lower than those observed in mice infected with either Delta or Omicron at 3dpi. Viral RNA was also detected in the medulla oblongata of mice infected by all these virus strains at 3dpi. At this time point, mice infected with the Delta virus display a marked upregulation of inflammatory makers both in the lungs and brains. Upregulation of inflammatory markers was also observed in the brains of mice infected with Omicron but not in mice infected with the Wild-type virus, suggesting that during the initial phase of the infection only the Delta and Omicron variants induce strong inflammatory response in the brain. At the humane endpoint/8dpi, mice infected by any of these strains display elevated levels of viral RNA and upregulation of a subset of inflammatory markers in the lungs. There was also a significant increase in viral RNA in the brains of mice infected with Wild-type and Delta, as compared to 3dpi. This was accompanied by an increase in the expression of most cytokines and chemokines. In contrast, mice infected with the Omicron variant showed low levels of viral RNA and downregulation of cytokines and chemokines expression at 8dpi, suggesting that brain inflammation by this variant is attenuated. Reduced RNA levels and downregulation of inflammatory markers was also observed in the medulla oblongata and olfactory bulbs of mice infected with Omicron, while infection by Wild-type and Delta resulted in high levels of viral RNA and increased expression of inflammatory makers in these organs.

## Introduction

Coronavirus Disease 2019 (COVID-19) is an infectious disease caused by the severe acute respiratory syndrome coronavirus 2 (SARS-CoV-2) that has affected millions of individuals worldwide. SARS-CoV-2 is a β-coronavirus with a positive-sense RNA genome very similar to SARS-CoV-1 (80%), but more infectious and transmissive due to a higher reproductive number (R0) [1]. SARS-CoV-2 contains five viral proteins, with nucleocapsid (N) protein coating and packaging for RNA genome, membrane (M) protein incorporating viral components into new virions, envelope (E) protein for viral assembly, and spike (S) protein participating in viral entry into host cells [2]. The early mutation found in SARS-CoV-2 is the Spike protein D614G mutation, which is also carried by many SARS-CoV-2 variants, including Delta (B.1.617.2) and Omicron (B.1.1.529) [3]. The D614G mutation results in increased infectivity by assembling more spike (S) protein on the virions and has enhanced viral loads in lung epithelial cells and the upper respiratory tract of COVID-19 patients. [4-5].

The Delta mutations have a concerning connection in causing more severe diseases. The Delta (B.1.617.2) variant can fuse the cellular membranes more efficiently, infect target cells much faster, and has a higher virion-releasing advantage over other SARS-CoV-2 variants. [6-7] Some of the spike mutations responsible for these changes are T478K and P681R. [8-9]. Different from other variants, Omicron (B.1.1.529) is found to be more dependent on the endocytic pathway for host cell entry [17]. Omicron variant has a large repertoire of mutations in the spike protein that can increase viral transmission and enhance cellular attachment [10-12]. However, studies have shown that the Omicron variant is compromised in the efficiency of viral fusion and viral replication competence [13-14], resulting in reduced viral load, attenuated inflammation, and overall severity of lung damage [15-16].

SARS-CoV-2 is a respiratory virus that can cause acute respiratory distress syndrome as well as neurological complications [18]. Patients being hospitalized due to acute infection have reported distinct neurological symptoms, including agitation, confusion, headache, and impaired consciousness [12, 19]. Severe neuronal damage was observed in the post-mortem brains from patients who died of SARS-CoV-2. Neurological damage included hemorrhage with infarction or white matter lesion, venous thrombosis [20-21] or thrombotic ischemic infarction [21-22], ischemic necrosis [23], edema [24], perivascular or microvascular congestion and injury [23], and hypoxic alterations [25-27]. Cerebrospinal fluid (CSF) samples collected from patients with moderate to severe COVID-19 showed elevated cytokine levels [28,29]. However, since very few clinical samples detected the presence of viral RNA and viral proteins [30], these neurological complications did not appear to be caused by direct viral infection in the brain [20-28]. Nevertheless, several laboratories have shown that SARS-CoV-2 can infect brain cells by crossing the blood-brain barrier and induce neuroinflammation [31]. Infection of human neural progenitor cells shows enhanced expression of viral transcription and metabolic processes, highlighting the ability of SARS-CoV-2 to hijack the host neuron cells to replicate [32]. SARS-CoV-2 can also infect non-neuronal cells, inducing upregulation of genes related to pro-inflammatory response and endoplasmic reticulum stress response (ER) [33]. It has also been reported that the S1 subunit of the S protein can cross the blood-brain barrier and activate microglial cells to increase cytokine release and inflammasome activities [34-36]. In this study, to compare the infectivity in the lung and brain of Wild-type, Delta, and Omicron in an animal model, we performed infections by these three strains of SARS-CoV-2 in the well-characterized K18-hACE2 transgenic mouse model.

## Materials and Methods

### Mice and viruses

All animal procedures were performed at the Hastings Foundation and the Wright Foundation BSL3 facility of the Keck School of Medicine at the University of Southern California (USC) and were approved by the Institutional Animal Care and Use Committee (IACUC) and the Institutional Biosafety Committee (IBC) of USC.

K18-hACE2 transgenic mice were obtained from the Jackson Laboratory (strain #: 034860). Female mice between the age of 8-10 weeks old were intranasally inoculated with SARS-CoV-2 and variants at 10^4 PFU and euthanized at set end point of 3dpi or the humane endpoint or 8 dpi which ever came first.

The Wild-type, Delta and Omicron SARS-CoV-2, all noted as variants of concern (VOC) by the United States’ Center for Disease Control and Prevention (CDC) were obtained from the BEI repository (Wild-type (USA-WA1/2020, catalog: NR-52281), Delta (B.1.617.2, catalog: NR-55671), and Omicron (B.1.1.529, catalog: NR-56461). All virions were propagated and tittered in Vero E6 cells overexpressing human ACE2 (VeroE6-hACE2) obtained from Dr. Jae Jung.

### Measurement of viral burden

Total RNA was collected from lungs, brains, olfactory bulbs and medulla oblongata of infected mouse tissues at the indicated time points using TRIzol reagent (Thermo Fisher Scientific) according to the manufacturer’s instructions. Complementary DNA (cDNA) was synthesized from DNase-treated RNA using iScript Reverse Transcription Supermix for RT-qPCR (BIO-RAD catalog: 1708841). 100ng of RNA template in a 10μl volume reaction was used with 2μl of iScript RT Supermix. Reaction protocol for cDNA synthesis was as follows: priming 5 min at 25°C, reverse transcription 20min at 46°C, RT inactivation 1min at 95°C. Copies of the SARS-CoV-2 nucleocapsid (N1) gene were determined using IDT Primetime Gene Expression Master mix (Cat#1055770). 2019-nCoV_N1 primer set was purchased from IDT (Cat#10007007). The primer sequence of N1 gene: Forward Primer: 5’-GAC CCC AAA ATC AGC GAA AT-3’ Reverse Primer: 5’-TCT GGT TAC TGC CAG TTG AAT CTG-3’ Probe: 5’-FAM-ACC CCG CAT TAC GTT TGG TGG ACC BHQ1-3’. Murine actin B (mActB) control primers were purchased from IDT (Mm.PT. 39a.22214843.g Cat# 1077852).10μl volume reaction was used with 5μl Primetime Gene Expression Master mix(2x), 1μl cDNA template, 0.75μl for N1 primer probe or 1μl for mAct B primer-probe (final reaction concentration: 500nM). Reaction protocol for RT-qPCR was as follows: Primer melting for 15 min at 50°C, Pre-denaturing for 2 min at 95°C, denaturation 3 sec at 95, annealing 30 sec at 55°C (40 cycles). Known genomic equivalent standards for SARS-CoV-2 quantification were run simultaneously.

### Measurement of cytokines and chemokines mRNA

RNAs from the lungs, brains, olfactory bulbs and medulla oblongata were extracted using the kit mentioned above. Complementary DNA (cDNA) was synthesized from DNase-treated RNA using iScript Reverse Transcription Supermix for RT-qPCR (BIO-RAD catalog: 1708841). 20μl volume reaction was used with 4ul of iScript RT Supermix and 200ng RNA template input. The protocol for cDNA synthesis was as follows: priming 5 min at 25°C, reverse transcription 20min at 46°C, RT inactivation 1min at 95°C. Cytokine and chemokine expression were determined using SsoAdvanced Universal SYBR Green Supermix (BIO-RAD catalog: 1725271). 10 μl volume reaction was used with 5 μl SsoAdvanced Universal SYBR Green Supermix(2x), 1 μl cDNA template, and 0.5 μl each forward and reverse primer (final reaction concentration: 500nM). Reaction protocol RT-qPCR (SYBR Green) was as follows: Initial DNA denature 30 sec at 95°C, denature 15 sec at 95°C, annealing/extension 30 sec at 60°C (40 cycles). The melt curve was run after the completion of qPCR to assess the production of single products. Primer sets were: CXCL9(F:CCTAGTGATAAGGAATGCACGATG;R:CTAGGCAGGTTTGATCTCCGTTC); CXCL10 (F: ATCATCCCTGCGAGCCTATCCT; R: GACCTTTTTTGGCTAAACGCTTTC);CCL8(F:GGGTGCTGAAAAGC-TACGAGAG;R: GGATCTCCATGTACTCACTGACC); VEGF-a (F: GCACTGGACCCTGGCTTTAC; R: ATCGGAC-GGCAGTAGCTTCG);CCL2(F:TGTTCACAGTTGCCGGCTG;R: GCACAGACCTCTCTCTTGAGC); IFN-γ (F: CAGCAACAGCAAGGCGAAAAAGG; R: TTTCCGCTTCCTGAGGCTGGAT);TNF-α(F:GGTGCC-TATGTCTCAGCCTCTT; R:GCCATAGAACTGATGAGAGGGAG);CXCL11(F:CCGAG-TAAGGCTGCGACAAAG; R:CCTGCATTATGAGGCGAGCTTG); IL-6 (F: ACCCCAATTTCCAATGCTCTCCT; R: ACGCACTAGGTTTGCCGAGTA); IL-1β (F: TGGACCTTCCAGGATGAGGACA; R: GTTCATCTCGGAGCCTG-TAGTG); IL-10 (F: CGGGAAGACAATAACTGCACCC; R: CGGTTAGCAGTATGTTGTCCAGC); GM-CSF (F: CCTGGGCATTGTGGTCTACAG; R: GGCATGTCATCCAGGAGGTT); G-CSF (F: GCAGACACAGTGCCTAA-GCCA; R: CATCCAGCTGAAGCAAGTCCA)

### Immunohistochemistry

Lung and brain tissue from mice were fixed in 10% formalin, embedded in paraffin, and sectioned (4–5 μm thickness). Tissue slides were baked at 60°Cfor 30 min, deparaffinized with xylene and rehydrated using ethanol. Slides were then boiled for 20 min in antigen retrieval buffer (Tris-EDTA buffer, pH9.0), and non-specific binding sites were blocked using 0.3% H_2_O_2_ as well as protein blocking reagent for 5min. After blocking, the sections were incubated with SARS-CoV-2 antibody against the nucleocapsid (N) protein (ThermoFisher MA536086; 1:28,000 dilution) for 15min at room temperature. After washing, the slides were incubated with the secondary antibody (Abcam ab64261) for 8min. Next, the chromogen DAB was applied to the slides for 5 min and slides were counterstained using hematoxylin for 10 min [32]. The slides were viewed in the bright field at 4x and 10x magnification on a Keyence microscope.

## Results

### Disease progression of SARS-CoV-2 infection of K18-hACE2 mice

K18-ACE2 transgenic mice were infected with 10^4 PFU of either Wild-type, Delta (B.1.617.2) or Omicron (B.1.1.529) SARS-CoV-2 and their weight and health conditions were recorded over a period of up to 8 days. Mice infected with the Wild-type virus started to show weight loss at 6dpi and reached the humane endpoint (HEP) between 6dpi and 7dpi (Figure 1). Disease progression was slightly faster in mice infected with the Delta variant, which began to lose weight at 4dpi and reach humane endpoint as early as 5dpi. Weight loss and moderate to severe neurological signs (head tilting, poor ambulation) are the prominent criteria for the humane endpoint in mice infected with the Wild-type and Delta SARS-CoV-2. In contrast, mice infected with the Omicron variant do not display any weight loss up to or past 8dpi and invariably survive the infection with no sign of neurological symptoms.

**Figure 1.**
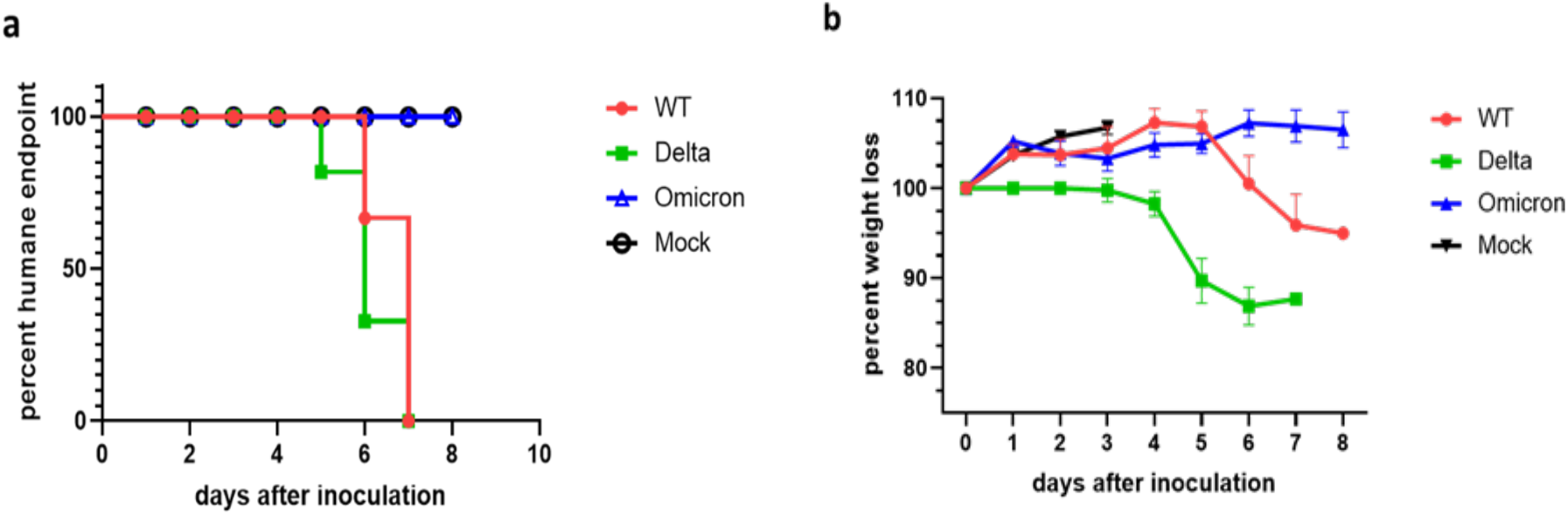
Probability of survival and body weight of SARS-CoV-2 infected K18-hACE2 mice. a) Mice were infected at 10^4 PFU with the indicated SARS-CoV-2 virus (Mock= uninfected; WT=Wild-type). The panel shows the percentage of mice that reached humane endpoint in days after inoculation (a.k.a. infection) (n=6). Mice infected with Omicron did not reach the humane endpoint and were euthanized at 8 days post infection (dpi). b) Weight loss of uninfected mice (mock) and mice infected with Wild-type, Delta, and Omicron (n=6). Mean ± s.e.m. Note: uninfected mice weight is only shown over 3 days.

### Pulmonary inflammation and infection of K18-hACE2 mice

Based on the differences in clinical manifestations between mice infected with the Wild-type, Delta, or Omicron SARS-CoV-2, we compare the viral load and inflammation in the lung at 3dpi versus humane endpoint. Mice infected with the Omicron variant were euthanized at day 8dpi to make a meaningful comparison with the mice infected with Wild-type or Delta. At 3dpi, the lungs of mice infected by the Wild-type or Delta SARS-CoV-2 show comparable viral loads (10^7^-10^8^ copy numbers/μg RNA), while mice infected with Omicron showed significantly lower levels of viral RNA (10^5^ copy numbers/μg RNA) (Figure 2a). Notably, infection by the Delta variant induced significantly higher levels of chemokine mRNAs such as CXCL9, CXCL10, CXCL11, and CCL2 than Wild-type and there was negligeable upregulation of cytokines and chemokines mRNA levels in mice infected with the Omicron at 3dpi. (Figure 2c). At the HEP, we detected higher levels of viral RNA in the lungs of mice infected with the Wild-type and Delta (>10^8^ copy numbers/μg RNA) than in the lungs of mice infected with Omicron (∼10^6^ copy numbers/μg RNA) at 8dpi (Figure 2b). Nevertheless, mice infected with Omicron display levels of inflammatory markers transcripts that were comparable to the levels observed in mice infected with Wild-type or Delta (Figure 2d). Thus, despite the lower levels of viral RNA, pulmonary inflammation is not attenuated in mice infected with Omicron.

**Figure 2.**
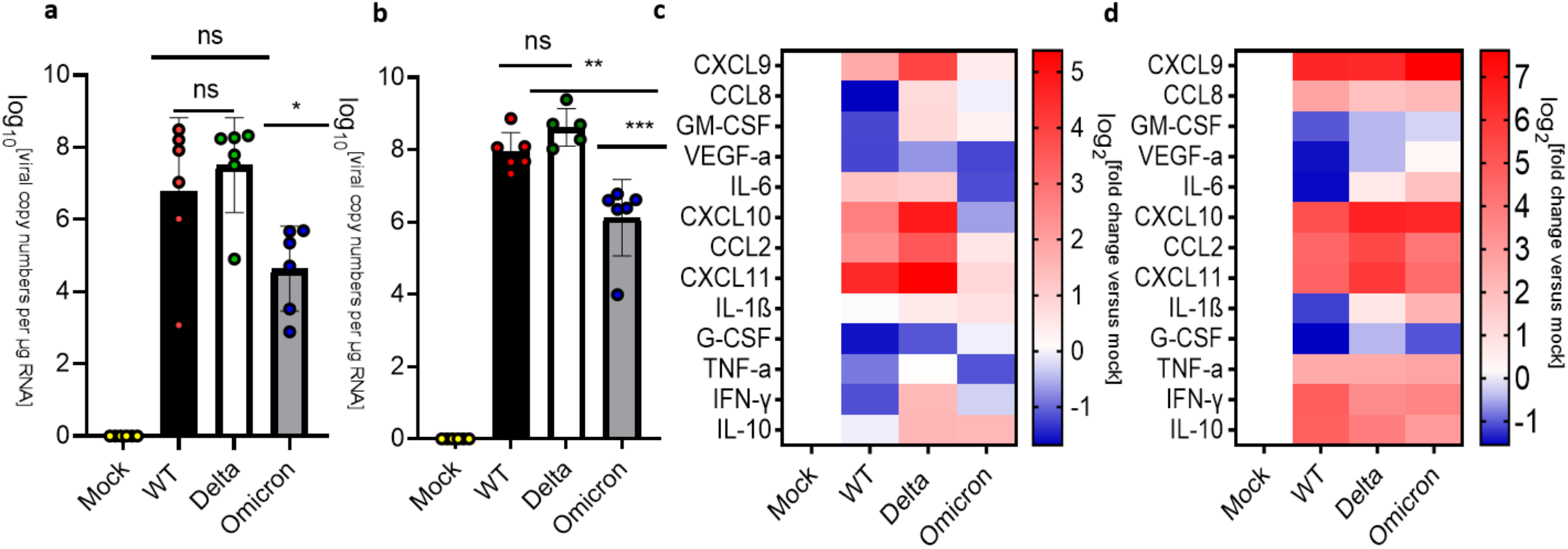
Analysis of viral RNA and cytokines/chemokines expression in the lungs of SARS-CoV-2 infected K18-hACE2. a) Uninfected mice (Mock) were compared against mice infected with SARS-CoV-2 Wild-type (WT), Delta, and Omicron at 10^4 PFU. Mice infected with Delta and Wild-type were euthanized at 3dpi or humane endpoint(5-7dpi). Mice infected with the Omicron variant were euthanized at 3dpi and 8dpi. a-b) qPCR of viral copy numbers in the lung at 3 dpi and humane endpoint/8dpi. Mean with SD. One-way ANOVA with Tukey’s test was performed *P< 0.05**P< 0.01, ***P< 0.001 a) qPCR of SARS-CoV-2 viral copy numbers at 3dpi. Mock (n=4), Wild-type, Delta, and Omicron (n=6) b) qPCR of SARS-CoV-2 viral copy numbers at the humane endpoint/8dpi. Wild-type (n=6), Delta (n=5), and Omicron (n=6). c-d) Heatmap of cytokines and chemokines levels in the lung. The graph was plotted with Mean. The fold change was calculated using the 2–ΔΔCt method and compared with mock-infected animals. The log2[fold change] was plotted in the corresponding heat map c) Heatmap of cytokines and chemokines levels in the lung at 3pi. Mock(n=4), Wild-type (n=6), Delta (n=6), Omicron (n=5). d) Heatmap of cytokines and chemokines levels in the lung at the humane endpoint/8dpi. Mock(n=4), Wild-type (n=6), Delta (n=5), Omicron (n=5).

### Infection and inflammatory markers in the brain of K18-hACE2 mice

COVID-19 can have profound effects on the brain and SARS-CoV-2 has been detected in the human brain at autopsy [52]. Numerous studies have examined the infection of the brain by SARS-CoV2 and suggested that the virus may enter the human brain through three main routes [53-54] (supplementary Figure 1). Here, we examined the brain of mice infected by Wild-type, Delta and Omicron. Note that the medulla oblongata was separated and analyzed independently of the rest of the brain (see Figure 3.4). Mice infected with the Wild-type virus did not show a significant amount of viral RNA in the brain at 3dpi. In contrast, the brains of mice infected with the Delta virus displayed relatively high levels of viral RNA at 3 dpi (∼10^5^ copy numbers/μg RNA). Interestingly, the brains of mice infected with Omicron also show higher levels of viral RNA than mice infected with Wild-type virus at 3dpi (>10^3^ copy numbers/μg RNA) (Figure 3a). The analysis of cytokines and chemokines mRNAs showed that the brains of mice infected with Omicron display upregulation of the inflammatory markers, while the brains of mice infected with Delta showed increased levels only for a subset of these markers, despite the high level of viral RNA at 3dpi (Figure 3c). The brains of mice infected with the wild-type virus only showed upregulation of TNF-alpha at 3dpi. At the HEP, mice infected with Wild-type and Delta showed high levels of viral RNA (10^8^-10^10^ copy numbers/μg RNA) (Figure 3b). Mice infected with Wild-type or Delta also showed a significant upregulation of inflammatory markers (Figure 3c). In contrast, mice infected with Omicron showed low levels of viral RNA (∼10^2^-10^3^copy numbers/μg RNA) and downregulation of inflammatory markers at 8dpi (Figure 3d). In agreement with these results, we observed a wide distribution of SARS-Co-V2 nucleocapsid (N) protein in the brains of mice infected with Wild-type and Delta at the HEP, while mice infected with Omicron had no detectable virus in the brain at 8dpi (Figure 3 e-h; supplementary Figure 2). Collectively, these data demonstrate differential levels of infection and inflammatory responses in mice infected by Wild-type, Delta and Omicron.

**Figure 3.**
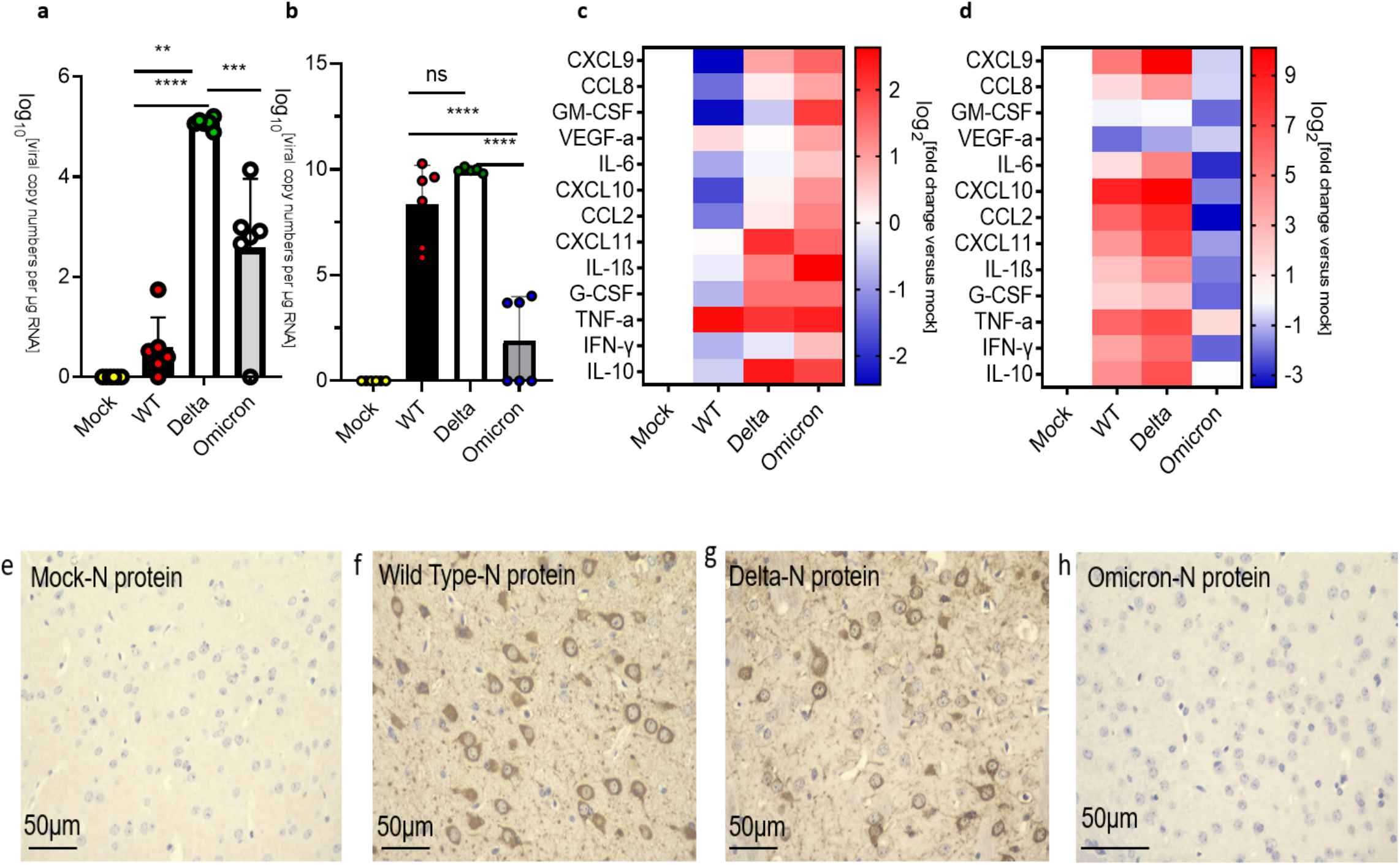
Analysis of viral RNA, viral protein, and cytokines/chemokines expression in the brains of SARS-CoV-2 infected K18-hACE2 mice. a-d) Uninfected mice (Mock) were compared against mice infected with SARS-CoV-2 Wild-type (WT), Delta, and Omicron at 10^4 PFU. Mice infected with Wild-type and Delta were euthanized at 3dpi and humane endpoint (5-7dpi), and mice infected with the Omicron variant were euthanized at 3dpi and 8dpi. a-b) qPCR of viral copy numbers in the brain at 3 dpi (a) and humane endpoint/8dpi (b). Mean with SD. One-way ANOVA with Tukey’s test was performed. **p<0.01, ***p<0.001, ****P< 0.0001 a) qPCR of SARS-CoV-2 viral copy numbers at 3dpi. Mock (n=4), Wild-type, Delta, and Omicron (n=6) b) qPCR of SARS-CoV-2 viral copy numbers at the humane endpoint/8dpi. Mock (n=4), Wild-type (n=6), Delta(n=5) and Omicron (n=6). c-d) Heatmap of cytokines and chemokines levels. The graph was plotted with Mean. The fold change was calculated using the 2–ΔΔCt method and compared with mock-infected animals. The log2[fold change] was plotted in the corresponding heat map c) Heatmap of cytokines and chemokines levels in the brain at 3dpi. Mock(n=4), Wild-type, Delta, and Omicron (n=6). d) Heatmap of cytokines and chemokines levels in the brain at the humane endpoint/8dpi. Mock(n=4), Wild-type (n=6), Delta (n=5), Omicron (n=5). e) Representative images showing IHC staining for the nucleocapsid (N) protein of SARS-Co-V2 in the brain of mice infected with Wild-type, Delta and Omicron at HEP/8dpi. e-h) Immunohistochemical (IHC) staining of brains of mice infected with Wild-type (f), Delta (g) and Omicron (h). Mock infected is shown in (e). The photomicrographs shown are representative of the images obtained from mice infected by each SARS-Co-V2 strain. Bars: 50 μm.

**Figure 4.**
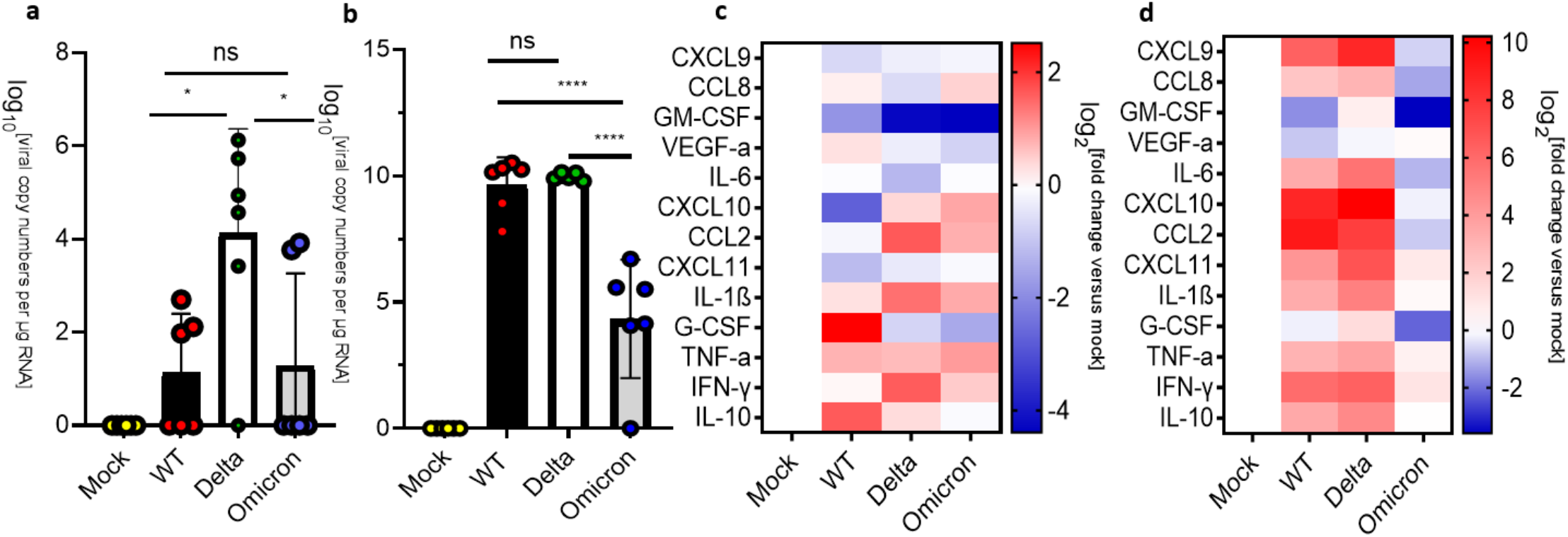
Analysis of viral RNA and inflammatory markers in the medulla oblongata of SARS-CoV-2 infected K18-hACE2 mice. Uninfected mice (Mock) were compared against mice infected with SARS-CoV-2 Wild-type (WT), Delta, and Omicron at 10^4 PFU. Mice infected with Delta and Wild-type were euthanized 3dpi or at the humane endpoint (5-7dpi). Mice infected with the Omicron variant were euthanized at 3dpi and 8dpi. a) qPCR of viral copy numbers at 3dpi. Mean with SD. One-way ANOVA with Tukey’s test was performed ****P< 0.0001 Mock (n=4), Wild-type (n=6), Delta(n=5) and Omicron (n=6) b) qPCR of viral copy numbers at humane endpoint/8dpi. Mean with SD. One-way ANOVA with Tukey’s test was performed *P< 0.01 Mock (n=4), Wild-type, Delta, and Omicron (n=6) c) Heatmap of cytokines and chemokines levels at 3dpi. Mock (n=4), Wild-type, Delta, and Omicron (n=6). d) Heatmap of cytokines and chemokines levels at the humane endpoint/8dpi. The graph was plotted with Mean. The fold change was calculated using the 2–ΔΔCt method and compared with mock-infected animals. The log2[fold change] was plotted in the corresponding heat map. Mock(n=4), Wild-type (n=6), Delta (n=5), Omicron (n=6).

### Infection and inflammation makers in the medulla oblongata of K18-hACE 2 mice

Since the human medulla oblongata has high levels of ACE2 expression [49], it represents a possible route for SARS-CoV-2 infection of the brain. We examined the medulla oblongata of mice infected with these SARS-CoV-2 strains at 3dpi and observed significantly higher levels of viral RNA in mice infected with Delta than either the Wild-type or Omicron (Figure 4a). However, infection by any of these SARS-CoV-2 strains resulted in mild upregulation of inflammatory markers including IL-1β, TNF-α, and IL-10 at 3dpi (Figure 4c). At HEP, we observed a major increase in the levels of viral RNA in the medulla oblongata of mice infected with Wild-type or the Delta variant (∼10^10^ copy numbers/μg RNA), while a less severe increase in viral RNA levels was observed in the medulla oblongata of mice infected with the Omicron variant (∼10^4^ copy numbers/μg RNA) at 8dpi (Figure 4b). At the HEP, inflammatory markers were significantly upregulated in mice infected with Wild-type or Delta but downregulated in mice infected with the Omicron variant at 8dpi (Figure 4 d).

### Infection and inflammation markers in the olfactory bulbs of K18-hACE 2 mice

The loss of smell observed in individuals infected by SARS-CoV-2 suggests that the olfactory bulb may be an important site of SARS-CoV-2 infection [50-51]. We therefore examined the levels of viral RNA and inflammatory makers in the olfactory bulbs of mice infected with Wild-type, Delta, and Omicron at the HEP/8dpi. As seen in the medulla oblongata, high levels of viral RNA (∼10^9^ copy numbers/μg RNA) and upregulation of inflammatory markers were only seen in in the olfactory bulbs of mice infected with Wild-type and Delta. The olfactory bulbs of mice infected with Omicron showed lower levels of viral RNA (∼10^5^ copy numbers/μg RNA) and downregulated inflammatory markers (Figure 5a,5b).

**Figure 5.**
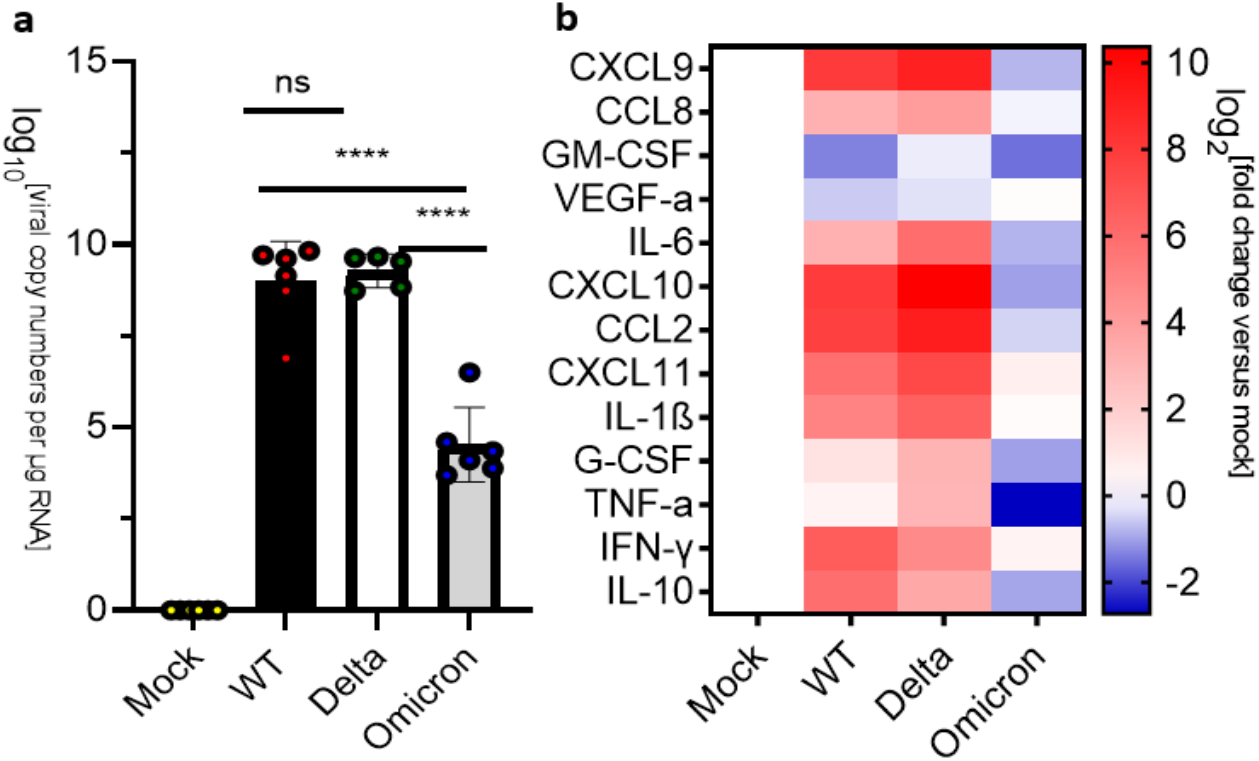
Analysis of viral RNA and inflammatory markers in the olfactory bulbs of SARS-CoV-2 infected K18-hACE2 mice at HEP/8dpi. Uninfected mice (Mock) were compared against mice infected with SARS-CoV-2 Wild-type (WT), Delta, and Omicron at 10^4 PFU. Mice infected with Wild-type or Delta were euthanized at the humane endpoint(5-7dpi), while mice infected with the Omicron variant were euthanized at 8dpi. a) qPCR of viral copy numbers in the OB at the humane endpoint/8dpi. Mean with SD. One-way ANOVA with Tukey’s test was performed ****P< 0.0001 Mock (n=4), Wild-type (n=6), Delta(n=5) and Omicron (n=6) b) Heatmap of cytokines and chemokines levels at the humane endpoint/8dpi. The graph was plotted with Mean. The fold change was calculated using the 2– ΔΔCt method and compared with mock-infected animals. The log2[fold change] was plotted in the corresponding heat map. Mock(n=4), Wild-type (n=6), Delta (n=5), Omicron (n=6). Note: olfactory bulbs were not analyzed at 3dpi.

## Discussion

The neurological impact of SARS-CoV-2 has been debated since the start of the COVID-19 pandemic. Studies on the relationship between viral loads and neurological inflammation and the impact of different SARS-CoV-2 variants on the nervous system have been limited. Here we compared the infectivity by Wild-type, Delta, and Omicron SARS-CoV-2 in transgenic K18-hACE2 mice and characterized the progression of infection and inflammatory response in the lung and brain of these animals. We observed that transgenic mice infected with Omicron display minimal weight loss and reduced disease progression as compared to mice infected with the Wild-type and Delta variant, which is in line with infection studies in hamsters [37-39]. Mice infected with the Delta variant have higher viral RNA levels and high levels of expression of inflammatory markers levels at the humane endpoint in the lung, suggesting that this SARS-CoV-2 variant results in enhanced lung infection and cellular damage. The high levels of lung inflammation caused by the Delta variant were also observed in other studies using the K18-hACE2 mice model [40-41]. Interestingly, although viral RNA levels in the lung of Omicron-infected mice are significantly lower, infection by this variant resulted in robust upregulation of inflammatory markers at both 3dpi and 8dpi as compared to Wild-type and Delta at 3dpi and HEP. As increased expression of proinflammatory cytokines has been reported to be related to reduced lung function and tissue damage in both acute and post-acute infections [40-42], the high expression level of cytokines/chemokines genes we observed in the lungs of mice infected by either Wild-type, Delta, or Omicron is likely to cause lung tissue damage and reduced lung function in post-acute infections. Many studies have shown significant and long-lasting neurological manifestations of SARS-CoV-2 infection [53]. Our analysis of the brains of mice infected with the Omicron variant shows reduced viral RNA levels as compared to the brains of mice infected with Wild-type or Delta at the humane endpoint/8dpi. We did not examine how the virus reaches the brain and whether it directly infects neurons or other brain cell types. However, studies by other groups have provided evidence of viral entry into the brain via distinct ways as well as direct infection of neurons in K19-hACE2 mice [55-57]. Regardless of the mechanism of entry, our study demonstrates that infection by different SARS-CoV-2 strains elicits distinct transcriptional responses of inflammatory markers in the brain of k18-hACE2. The brains of mice infected with Omicron show a significant upregulation of cytokines and chemokines at 3dpi. This increase appears to be transient, as these markers are all downregulated at 8dpi, suggesting that infection by this variant results in attenuated neurological consequences compared to mice infected with either Wild-type or Delta. This outcome agrees with the reported lack of neurological symptoms after the Omicron outbreak in human cohort studies [43-44].

We observed viral RNA and mild upregulation of inflammatory markers in the medulla oblongata of mice infected with Wild-type, Delta, and Omicron at 3dpi. This is consistent with the detection of the virus and activation of immune response cells in the brainstems of COVID-19 patients [45]. However, while the high levels of viral RNA and a significant upregulation of inflammatory genes were observed in mice infected with Wild-type and Delta, mice infected with the Omicron variant showed lower levels of viral RNA and downregulation of inflammatory genes at 8dpi. Lower levels of viral RNA and downregulation of inflammatory markers were also seen in the olfactory bulb of mice infected with Omicron at 8dpi as compared to Delta and Omicron at the human endpoint, a result that is similar to the low viral load observed in the olfactory bulb of hamsters infected with Omicron [46]. Moreover, in contrast to mice infected with Wild-type or Delta, the infection by Omicron did not result in any upregulation of inflammatory markers in the olfactory bulbs. These findings align with cohort studies that reported limited loss of smell and taste in patients infected with the Omicron variant [47-48]. Collectively, these data demonstrate that K18-hACE2 mice infected with Wild-type, Delta, and Omicron show distinct levels of viral RNA and inflammatory responses in the lung, brain, medulla oblongata and olfactory bulbs during the infection. Importantly, the inflammatory response in mice infected by the Omicron variant is limited to the early phase of infection and does not lead to health conditions that require euthanasia.

### Limitations of the study

The study has potential limitations. The transgenic K18-hACE2 mouse model is highly susceptible to lethality of SARS-CoV-2 infection and does not fully recapitulate infection in humans. This model is therefore not suitable to study long-term effects of SARS-CoV-2 infection (long COVID). Nevertheless, our data show that K18-hACE2 mice infected with the Omicron recover from the infection, suggesting that infection of K18-hACE2 mice by this variant may offer a model system to study long COVID. Another limitation of the study is in the use of RT-qPCR to measure changes in cytokines and chemokines mRNAs. While these transcriptional events are likely associated with changes in protein levels, we have not performed experiments to assess these changes.

## Acknowledgments

This work was performed at the Hastings Foundation and Wright Foundation Biosafety level 3 (BLS3) laboratory of USC and was funded by a research grant from the W.M. Keck Foundation to L.C. We thank the Pathology Core lab at USC for support with the immunohistochemical experiments.

## Author Contributions

Conceptualization, Y.H. and L.C.; methodology, Y.H, P.S, J.H.; formal analysis, J.H., P.S., J.H.; investigation, Y.H, P.S, J.H.; resources, J.H., L.C.; data curation, Y.H.; writing—original draft preparation, Y.H.; writing—review and editing, Y.H, J.H, L.C.; supervision, J.H., L.C.; project administration, J.H.; funding acquisition, L.C. All authors have read and agreed to the published version of the manuscript.

## Institutional Review Board Statement

The study was conducted in accordance with the Declaration of Helsinki, and approved by the Institutional Review Board of The University of Southern California. The animal study protocol was approved by the Institutional Animal Care and Use Committee of the University of Southern California.

## Data Availability Statement

Data are available from corresponding authors upon request.

