## Supplementary Figures 1 and 2 for "Differential outcomes of infection by wild-type SARS-CoV-2 and the B.1.617.2 and B.1.1.529 variants of concern in K18-hACE2 transgenic mice"

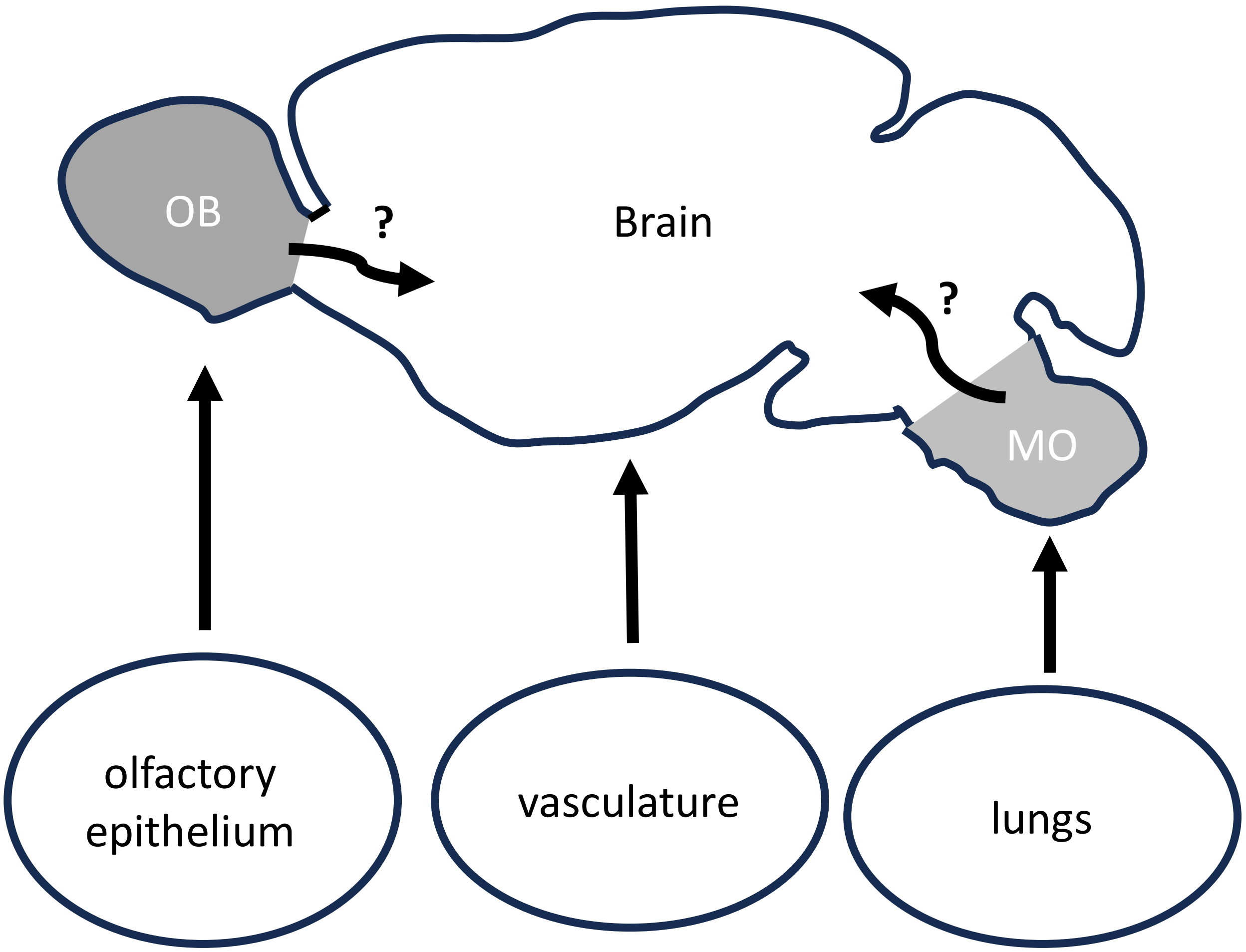


Supplementary Figure 1: Schematic representation of mouse brain with possible routes of CNS infection by SARS-CoV-2. 1. Olfactory route: 2. Vasculature and crossing the blood-brain barrier; 3. Travels from the lungs to the medulla oblongata brain stem through the vagal nerve.  OB: olfactory bulb; MO: medulla oblongata

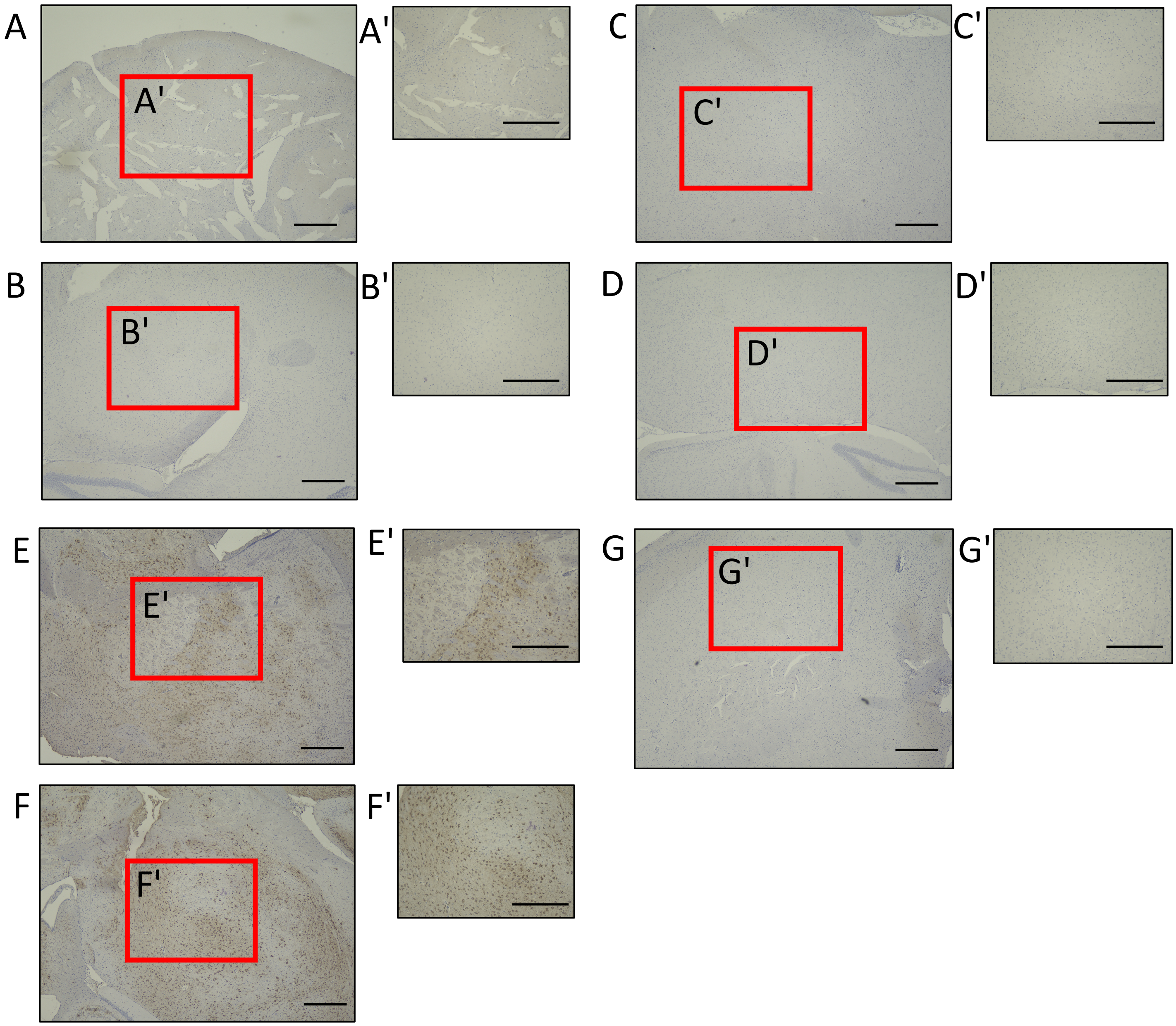


Supplementary Figure 2: Representative images showing IHC staining for the nucleocapsid (N) protein of SARS-Co-V2 in the brain of mice infected with Wild-type, Delta and Omicron at 3dpi and HEP/8dpi. Immunohistochemical (IHC) staining of brains of mice infected with Wild-type (B and E), Delta (C and F) and Omicron (D and G) at 3dpi (B-D), HEP (E-F) or 8dpi (G). Mock control brain is shown in A. The photomicrographs shown are representative of the images obtained from mice infected by each SARS-Co-V2 strain. A'-G' show higher magnification images of inset in panels A-G. Bars: 500 µm.
